# Size and composition of haplotype reference panels impact the accuracy of imputation from low-pass sequencing in cattle

**DOI:** 10.1101/2023.01.13.523894

**Authors:** A. Lloret-Villas, H. Pausch, A.S. Leonard

## Abstract

**Background:** Low-pass sequencing followed by sequence variant genotype imputation is an alternative to the routine microarray-based genotyping in cattle. However, the impact of haplotype reference panel composition and its interplay with the coverage of low-pass whole-genome sequencing data has not been sufficiently explored in typical livestock settings where only a small number of reference samples are available.

**Methods:** Sequence variant genotyping accuracy was compared between two variant callers, GATK and DeepVariant, in 50 Brown Swiss cattle with sequencing coverages ranging from 4 to 63-fold. Haplotype reference panels of varying sizes and composition were built with DeepVariant considering 501 cattle from nine breeds. High coverage sequencing data of 24 Brown Swiss cattle was downsampled to between 0.01- and 4-fold coverage to mimic low-pass sequencing. GLIMPSE was used to infer sequence variant genotypes from the low-pass sequencing data using different haplotype reference panels. The accuracy of the sequence variant genotypes imputed inferred from low-pass sequencing data was compared with sequence variant genotypes called from high-coverage data.

**Results:** DeepVariant was used to establish bovine haplotype reference panels because it outperformed GATK in all evaluations. Same-breed haplotype reference panels were better suited to impute sequence variant genotypes from low-pass sequencing than equally-sized multibreed haplotype reference panels for all target sample coverages and allele frequencies. F1 scores greater than 0.9, implying high harmonic means of recall and precision of called genotypes, were achieved with 0.25-fold sequencing coverage when large breed-specific haplotype reference panels (n = 150) were used. In absence of such large same-breed haplotype panels, variant genotyping accuracy from low-pass sequencing could be increased either by adding non-related samples to the haplotype reference panel or by increasing the coverage of the low-pass sequencing data. Sequence variant genotyping from low pass sequencing was substantially less accurate when the reference panel lacks individuals from the target breed.

**Conclusions:** Variant genotyping is more accurate with Deep-Variant than GATK. DeepVariant is therefore suitable to establish bovine haplotype reference panels. Medium-sized breed-specific haplotype reference panels and large multibreed haplotype reference panels enable accurate imputation of low-pass sequencing data in a typical cattle breed.

## Introduction

More than a million cattle are genotyped every year with microarray technology for the purpose of genomic prediction (1). Access to whole genome sequence variants can improve the accuracy of genomic predictions and facilitates the monitoring of trait-associated alleles (2). However, costs are still too high to sequence all individuals from a population to a sufficient coverage to call variants.

Low-coverage whole-genome sequencing (lcWGS) followed by genotype imputation has emerged as an alternative with comparable costs to genotyping microarrays but with substantially higher marker density (tens of millions versus tens of thousands) for obtaining genotypes for a target population (3–6). Sequencing coverage as low as 0.1-fold can be used to infer sequence variant genotypes that are as accurate as those obtained from genotyping microarrays, especially for rare variants, while sequencing coverage greater than 1-fold can have much higher accuracy (5). For many imputation methods, reference panels that are representative for the target populations are a prerequisite for the accurate imputation of genotypes from lcWGS (7–9). The 1000 Genomes Project (1KGP) and the Haplotype Reference Consortium (HRC) established such reference panels for several human ancestry populations (10, 11) and made them available through dedicated imputation servers (12). A bovine imputation reference panel established by the 1000 Bull Genomes project is frequently used to infer sequence variant genotypes for large cohorts of genotyped taurine cattle, thus enabling powerful genome-wide analyses at the nucleotide level (13). Sequenced reference panels are available for other animal species (14, 15). However, these haplotype panels lack diversity as they were established mainly with data from mainstream breeds and thus are depleted for individuals from local or rare populations.

An exhaustive set of variants and accurate genotypes are crucial to compile informative haplotype reference panels. The Genome Analysis Toolkit (GATK) has been frequently applied to discover and genotype sequence variants in large reference populations of many livestock species (3, 14). DeepVariant has recently emerged as an alternative machine learning-based variant caller (16). Several studies suggest that DeepVariant has superior genotyping accuracy over GATK (17–20). However, DeepVariant had rarely been applied to call variants in species other than humans (21, 22).

In this study, we benchmark sequence variant genotyping of DeepVariant and GATK in a livestock population. We then build haplotype reference panels of varying sizes and composition with DeepVariant, and use GLIMPSE to impute sequence variant genotypes for cattle that had been sequenced at between 0.01- and 4-fold genome coverage. We show that within-breed haplotype reference panels outperform multi-breed reference panels across all tested scenarios, provided that enough sequenced samples are available.

## Materials and methods

### Data availability and code reproducibility

Short paired-end whole-genome sequencing reads of 501 cattle from nine breeds were used: 327 Brown Swiss (BSW), 50 Fleckvieh, 13 Hereford, 57 Holstein, 2 Nordic Red, 14 Rätisches Grauvieh, 10 Simmental, 25 Tyrolean Grauvieh and 3 Wagyu cattle. Accession numbers for the raw data are available in the Supplementary file 1. Computational workflows were implemented using Snakemake (23) (version 7.5.0 or newer). The R software environment (version 4.0.2) and ggplot2 package (24) (version 3.3.2) were used to create figures and perform statistical analyses. Scripts and workflows are available online.

### Alignment, mapping quality and depth of coverage

Raw short sequencing reads were filtered with fastp (25) (version 0.23.1), and MultiQC (26) (version 1.11) was applied to collect the quality metrics across samples. Reads were split per read groups with gdc-fastq-splitter (27) (version 1.0.) and subsequently aligned with bwa-mem2 (28) using the *-M* and *-R* flags to a manually curated version of the current bovine Hereford-based reference genome (ARS-UCD1.2) (29) that included a Y chromosome as described in (30). Samblaster (31) (version 0.1.26), Sambamba (32),samtools (33, 34) (version 1.12), and Picard tools (35) (version 2.25.7) were used to deduplicate and sort the BAM files. We calculated average coverage with mosdepth (36) (version 0.3.2) considering all aligned reads that had MQ ≥ 10.

### Comparison of variant callers

#### Testing set

50 BSW cattle with coverages ranging from 4 to 63-fold were selected as testing set for a comparison between GATK and DeepVariant.

#### GATK

We used the BaseRecalibrator module of GATK (37, 38) (version 4.2.2.0) to adjust the base quality scores of the deduplicated bam files using 1 15,815,224 u nique positions from the Bovine dbSNP version 150 as known variants. Multi-sample variant calling was performed with the GATK HaplotypeCaller, GenomicsDBImport and GenotypeGVCFs modules according to the best practice guidelines (39, 40). We applied the VariantFiltration module for site-level filtration with thresholds indicated in (30) to retain high-quality SNP and INDELs.

#### DeepVariant + Glnexus

DeepVariant (16) (version 1.2) was run on the deduplicated bam files using the *WGS* Illumina-trained model, producing gVCF output per sample. The gVCF files were then merged and filtered using GLnexus (41) (version 1.4.1) with the *DeepVariantWGS* configuration but with the *revise_genotypes* flag set to false.

#### VCF imputation and statistics

We used Beagle 4.1 (42) (27Jan18.7e1) to improve genotype calls and impute sporadically missing genotypes from genotype likelihoods (*gl* mode). INDELs were left-normalised using bcftools (34) (version 1.12 or 1.15) *norm*. Variant and genotype counts, and Ti:Tv ratios were calculated with bcftools *stats* and bcftools *query*. VCF files were indexed with tabix (43, 44).

#### Variant annotation

Functional consequences of SNPs were predicted based on the Ensembl (release 104) annotation of the bovine reference assembly using the Variant Effect Predictor tool (VEP) (45) (version 106) with default parameter settings.

#### Variant accuracy evaluation

Microarray-derived genotypes from 33 cattle that also had sequence-derived genotypes (Supplementary File 1) were our truth chip set. We intersected the truth (microarray) and query (WGS variants) VCF files using bcftools *isec* with both the *-c none* (exact – only matching REF:ALT alleles are allowed) and *-c all* (position– all coordinate matches are allowed) flags, and retained biallelic SNPs with bcftools *view* to compare the genotypes. Three-way intersection overlaps were counted with bedtools *multiinter* (46) and visualised with UpSetR (47, 48). Since the microarray data contains fewer sites than WGS, we intersected the truth and query sets. Only positions where the truth genotypes were not homozygous for the reference allele (*i*.*e*., the variants that segregate within the target samples) were retained. We calculated recall (percentage of true positives in the query set), precision (proportion of matching genotypes in both truth and query sets), and F1 scores (harmonic mean of precision and recall) using hap.py (49) (version 0.3.9) on a per-sample basis. Agreement between the imputed variant alleles/genotypes and raw sequencing reads was assessed with Merfin’s k-mer-based filtering method (50) (commit fc4f89a). A k-mer database was prepared using Meryl (commit 51fad4b) with a k-mer size of 21 and minimum k-mer occurrence of 2 in the short sequencing reads. Variants that were poorly supported, *i*.*e*., the alternate sequence (variant and flanking regions) appeared less often in k-mers than the reference sequence did in a genotype-aware proportion, were filtered out.

We assessed Mendelian consistency in filtered but not-imputed data from parent-offspring pairs and trios (Supplementary File 2) using the bcftools *+mendelian* plugin (34). We calculated discrepancy rate as the number of inconsistent sites divided by the total number of non-missing sites. For duos (dam-offspring or sire-offspring) only homozygous sites were considered. When only one parent was available (duos), assessing discrepancy was only possible when the parent genotype was homozygous (0/0 or 1/1).

### Imputation of low-pass sequencing data

#### Haplotype panel generation

The BSW reference panels contained 150, 75 and 30 samples that were randomly selected from 303 BSW samples. The non-BSW panels contained 150, 75 and 30 samples that were randomly selected from 174 non-BSW samples. The multibreed panels were randomly selected from a combination of the above, and they contained 150 samples of which 50%, 25%, and 10% were BSW samples and the remaining were non-BSW. Three random replicates for each panel were created. A subset of 2,078 taurine samples of the 1000 Bull Genomes project (13) was used to generate a benchmark haplotype reference panel. Sequence variant genotypes were called for each panel with DeepVariant and sporadically missing genotypes were imputed with Beagle 4.1 (42) (27Jan18.7e1) as described above.

#### Truth sequencing set, truth variants and subsampling

Variants were called with DeepVariant and GLnexus as described previously for 24 BSW samples with coverage above 20-fold to generate a truth set for assessing imputation accuracy. The raw whole-genome sequencing reads of the 24 BSW samples were then downsampled with seqtk (51) to mimic 4x, 2x, 1x, 0.5x, 0.25x, 0.1x, and 0.01x coverage, and subsequently aligned to ARS-UCD12 as described previously.

Genotype likelihoods for the variants that are present in the haplotype reference panel were estimated from the subsampled read alignments with bcftools *mpileup* and bcftools *call*. These were then imputed using the different haplotype panels using GLIMPSE (52) (version 1.1.1). We used 2 Mb windows and 200 Kb buffer sizes during the chunk step followed by phasing and ligation to produce the final imputed variant calls.

#### Comparison of true and imputed variants

The accuracy of the imputed sequence variant genotypes was assessed with hap.py as described above. The minor allele frequency (MAF) of the imputed sequence variants was calculated with PLINK (53) (version 1.9). The estimated imputation quality was retrieved from the INFO flag from the VCF files produced by GLIMPSE with bcftools *query*. Pearson squared correlation between expected and actual dosages (*r*^2^) was calculated with the bcftools *stats*.

## Results

### Variant calling with GATK and DeepVariant

We compared sequence variant discovery and genotyping between GATK and DeepVariant in 50 Brown Swiss (BSW) cattle that had between 4 and 63-fold sequencing depth (19.26 ± 11.09) along the autosomes. GATK and DeepVariant identified 18,654,649 and 18,748,114 variants, respectively, of which 7.79% and 8.38% were filtered out due to low quality (Table 1). There were 16,147,567 filtered variants identified by both callers, but 1,053,716 and 1,292,671 variants were private to GATK and DeepVariant, respectively (Figure 1A). Overall, DeepVariant had more private SNPs than GATK, but GATK had more private INDELs than Deep-Variant (Supplementary Table 1). 416,642 variants had the same coordinates but different alternative alleles. These discrepant sites were primarily INDELs (83%, as opposed to the 12% of INDELs in all shared variants). Multiallelic sites accounted for 3.44% and 3.31% of the variants (0.33% and 0.28% of the SNPs, and 23.22% and 23.94% of the INDELs) that passed the quality filters of GATK and DeepVariant, respectively. Multiallelic sites were enriched among the variants private to either GATK or DeepVariant (Supplementary Table 2).

**Fig. 1.**
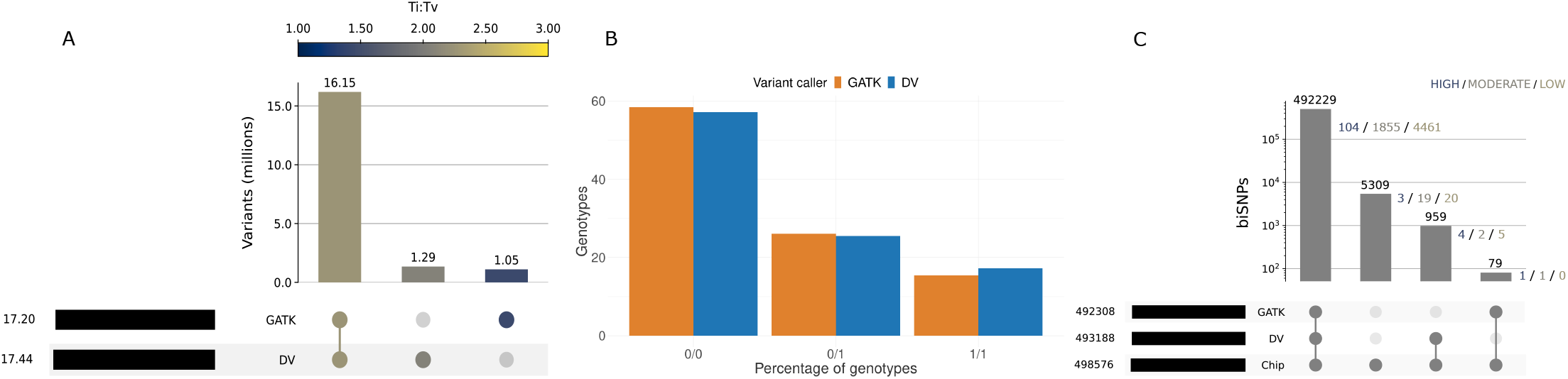
Variant call comparison between DeepVariant (DV) and GATK. a) Intersection of variants called with each variant caller (or both) and the Ti:Tv ratio of the biallelic SNPs of each set. b) Percentage of imputed genotypes called by each variant caller. c) Intersection of variant calls with truth genotyping arrays, where only positions intersecting truth are retained. Low, moderate, and high predicted impact variants from the intersection sets are indicated.

**Table 1.**
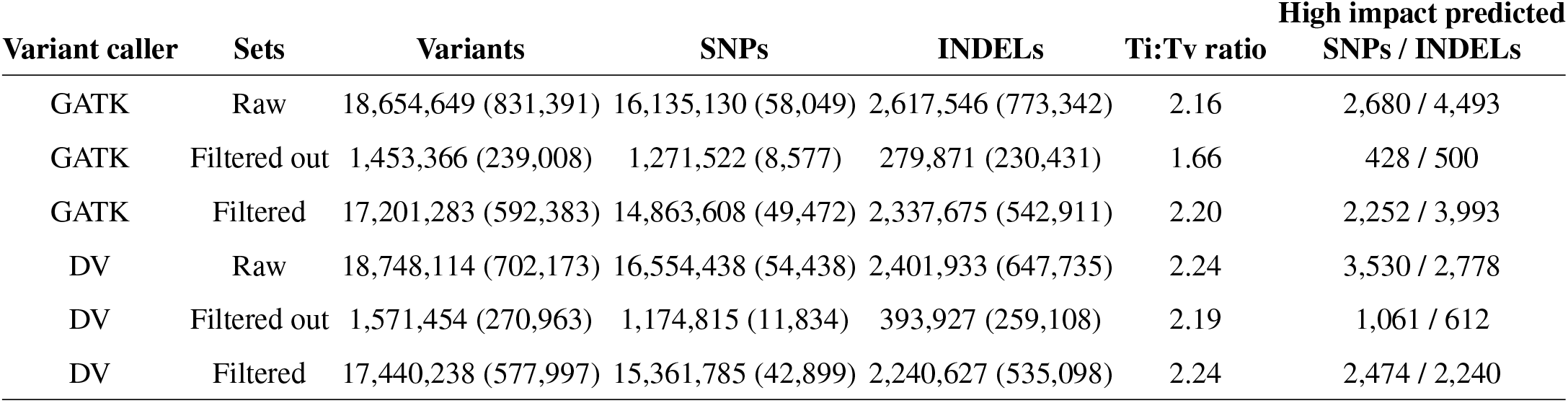
Summary of the variants called by GATK and DeepVariant (DV). Multiallelic sites are presented in parentheses. Ti:Tv ratios are restricted to biallelic SNPs. Functional consequences are predicted for biallelic SNPs / biallelic INDELs.

The biallelic variants called by GATK had a higher percentage of homozygous reference (HOMREF) and heterozygous (HET) genotypes whereas the biallelic variants called by DeepVariant had a higher percentage of homozygous alternative (HOMALT) genotypes (Figure 1B, Supplementary Figure 1A). Missing genotypes were very rare (*<*0.01%) for GATK-called biallelic variants but accounted for 2.72% of the DeepVariant-called genotypes (Supplementary Figure 1B). Beagle phasing and imputation increased the number of HET genotypes for both GATK - mainly transitioning from HOMREF - and DeepVariant - mainly due to the refinement of missing genotypes (Supplementary Figure 1C).

Functional consequences on the protein sequence were predicted for all biallelic variants. DeepVariant identified 9% more SNPs that were predicted to have a high impact on protein function than GATK (Table 1 & Supplementary Table 3). Around one fourth of the high impact SNPs detected by DeepVariant (24%) were not detected by GATK. GATK identified 78% more INDELs that were predicted to have a high impact on protein function than DeepVariant. More than half of the high impact INDELs detected by GATK (52%) were not detected by DeepVariant.

We investigated the ratio of transitions to transversions (Ti:Tv) to assess variant quality. Deviations from an expected genome-wide Ti:Tv ratio of ∼ 2.0-2.2 indicate random genotyping errors or sequencing artifacts (17, 20, 38, 54). The Ti:Tv ratio was 2.16 and 2.24 for raw SNPs identified by GATK and DeepVariant, respectively (Table 1). While the Ti:Tv ratio was higher (2.20) for the GATK variants that met the quality filters, variant filtration had no impact on the Ti:Tv ratio for SNPs called by DeepVariant. The Ti:Tv ratio of the filtered out SNPs was substantially lower for GATK (1.66) than DeepVariant (2.19). SNPs private to GATK had lower Ti:Tv ratios than the SNPs private to DeepVariant (Figure 1A). Substantial differences in the Ti:Tv ratio (0.81 points) existed between overlapping and GATK-private SNPs but were less (0.18 points) between overlapping and DeepVariant-private SNPs.

### Variant calling accuracy

Thirty-three sequenced cattle also had between 17,575 and 490,174 SNPs genotyped with microarrays. The filtered biallelic SNPs called with GATK and DeepVariant (query sets) were compared to those genotyped with the microarrays (truth chip set). The vast majority (98.82%) of the SNPs present in the truth chip set was called by both tools (Figure 1C). The overlap of SNPs present in the truth chip set was slightly higher for DeepVariant than GATK. 1.06% (n = 5,309) of the SNPs present in the truth chip set were not called by any of the software as biallelic SNPs. However, 3,497 of these SNPs were present at the same position but had different alternative alleles (*e*.*g*., multiallelic SNPs or INDELs) in DeepVariant/GATK while the other 1,812 positions were truly missing. Most of the biallelic SNPs private to the chip set (5,265) were also missing in the raw calls from the variant callers. DeepVariant filtered out more variants present in the truth chip set than GATK.

The analysis of variant effect predictions for the filtered variants revealed that most low/moderate/high impact variants were called by both GATK and DeepVariant (99.4%, 98.8%, and 92.8%, respectively). However, DeepVariant additionally called 5/2/4 biallelic SNPs predicted as low/moderate/high impact respectively, while GATK only called 0/1/1 (Figure 1C). Some of the low/moderate/high impact biallelic SNPs private to GATK (1 out of the 2) and DeepVariant (5 out of the 11) were called either as multiallelic SNPs or as INDELs by the other caller (Supplementary Table 4). Only half (1 out of 2) of GATK’s private variants annotated with low/moderate/high have minor allele frequencies (MAF) *>* 0.05, while most (9 out of 11) of DeepVariant’s do, suggesting that GATK misses more variants that might have larger impact in populations.

### Genotyping accuracy of variant calls

GATK and DeepVariant called 492,265 and 493,145 variants from the truth chip set, respectively. GATK missed (8.13%) and miscalled (10.13%) more truth variants than DeepVariant. Around 90.6% of the discrepancies between the sequence variant genotypes and the truth chip set in both variant callers were due to missing genotypes in the sequence set. Of those, GATK missed proportionally more HOMALT than DeepVariant and DeepVariant missed proportionally more HET variants. For the remaining *∼* 9.4% of mismatching genotypes (miscalled), also GATK miscalled proportionally more HOM variants and DeepVariant significantly miscalled proportionally more HET variants (Supplementary Figure 2). After imputation, however, the proportion of HET positions miscalled was higher in the GATK set and the proportion of HOMREF positions miscalled as HET was significantly higher in the DeepVariant set.

Recall, precision and F1 score of the filtered query sets were calculated to assess the genotyping accuracy for both variant callers. DeepVariant had strictly better F1 scores than GATK for the filtered data (mean of 0.9719 versus 0.9694, Figure 2A-B). The difference was small but significant (Wilcoxon signed-rank test, p=2.3*x*10^−10^). As expected, lower coverage (<20x) samples benefited from imputation, improving their F1 scores to be comparable to high coverage samples. Imputation improved GATK genotypes more than DeepVariant genotypes at lower coverages, potentially due to better calibration of genotype likelihoods, but DeepVariant was still strictly better above 7x coverage. Overall, DeepVariant still had a significantly higher mean F1 score for the imputed data (0.9912 versus 0.9907, Wilcoxon signed-rank test p=4.2*x*10^−05^, Figure 2C).

We further examined variant genotyping accuracy through Merfin (50). Merfin filters out variants when the proportion of “reference” and “alternate” k-mers for that variant from the sample’s short sequencing reads does not match the genotype and so is likely wrong. HET genotypes of both GATK and DeepVariant had less support from the sequencing reads, as they are harder to genotype correctly than HOM genotypes. For both HET and HOMALT, more of DeepVariant’s than GATK’s variants were supported (Figure 3A). The difference between the tools was statistically significant for both genotypes (two-sided paired Wilcoxon test, p_HET_=3.6*x*10^−19^, p_HOMALT_=1.8*x*10^−19^).

**Fig. 2.**
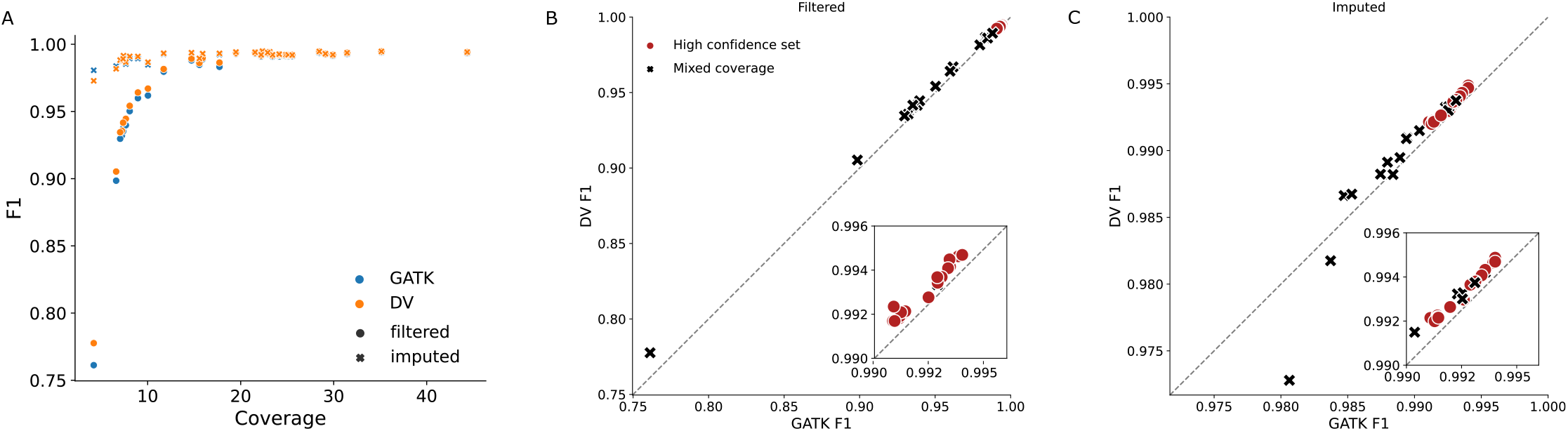
Comparison of the F1 values obtained with hap.py from GATK and DeepVariant (DV) variant calls against the truth chip set for 33 samples. a) Imputation improves genotype accuracy below 20x coverage but has minor impact above that. b) DV has a higher F1 score for every sample than GATK for post-filter variants. The high confidence set indicates the 17 microarray genotyped samples out of the 24 samples used later as a truth set for GLIMPSE imputation. c) Similar to (b) but for post-imputation variants.

**Fig. 3.**
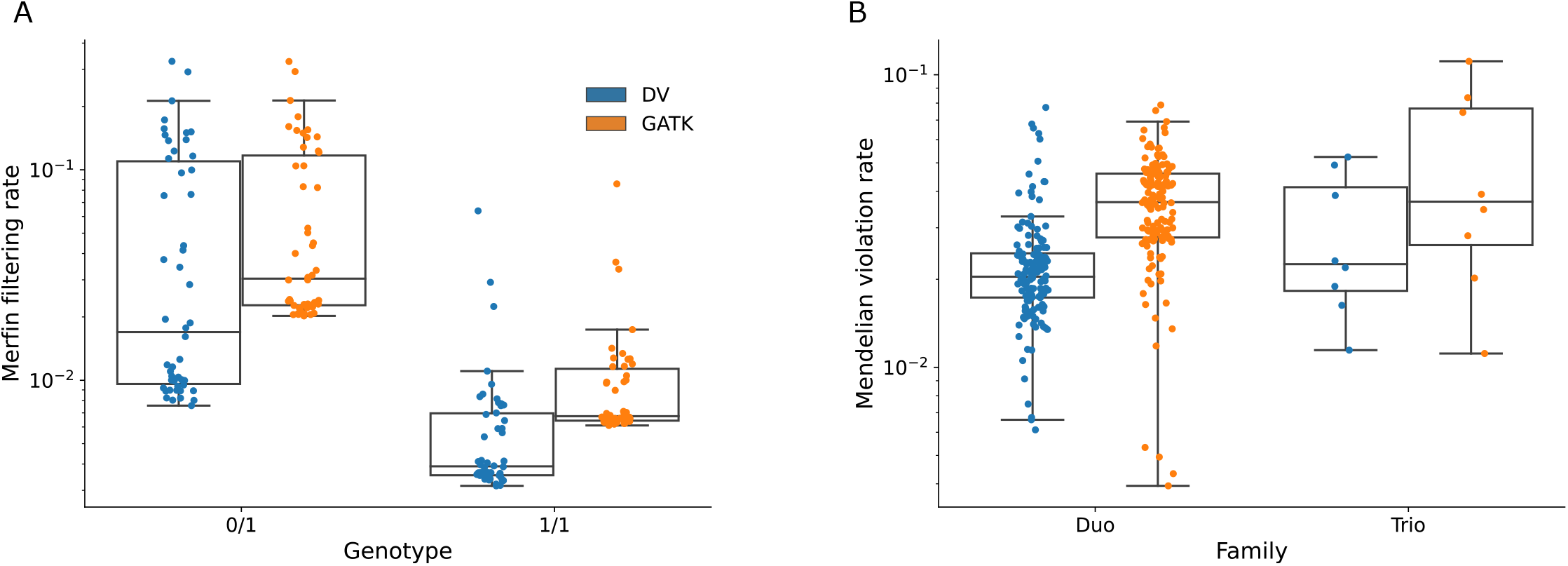
Genotyping accuracy of variant calls validated with sequencing reads and mendelian relationships. a) Filtering rate of heterozygous (0/1) and homozygous alternate (1/1) variant calls post-imputation for GATK and DV. Higher filtering rate indicates the genotype/allele is not consistent with k-mers from the same-sample sequencing reads. b)Mendelian violation rate for 206 separate samples, with either 2 family members (Duo) or all 3 (Trio). Mendelian violations are defined as genotypes in the offspring that could not have been inherited from the parents. In the case of duos, only homozygous variants can be assessed.

In addition, we compared Mendelian concordance rate among sequenced duos and trios across the two variant callers. There were only two family relationships in the previously examined 50 samples, and so we evaluated the concordance on a separate set of 206 samples (Supplementary File 2) forming 7 trios (both parents available) and 142 duos (one parent available). DeepVariant had less genotypes conflicting with Mendelian inheritance compared to GATK (2.3% versus 3.8%, Figure 3B, one-sided paired Wilcoxon signed-rank test p=1.3*x*10^−24^). This was due to DeepVariant calling both more genotypes that were compatible as well as fewer that were incompatible with parent-offspring relationship.

### Generation of a sequencing validation set for lcWGS imputation

We benchmarked the accuracy of low-pass sequence variant imputation in a target population consisting of 24 BSW samples with mean autosomal coverage of 28.12 ± 9.07-fold. DeepVariant identified 15,948,663 variants (87.77% SNPs and 12.23% INDELs) in this 24-samples cohort of which we considered 13,854,932 biallelic SNPs as truth set.

The sequencing reads of these 24 samples were randomly downsampled to mimic mid (4x and 2x), low (1x, 0.5x, 0.25x, and 0.1x), and ultralow (0.01x) sequencing coverage. We then aligned the reads to the reference sequence and produced genotype likelihoods from the pileup files. Subsequently, genotypes were imputed with GLIMPSE considering nine haplotype reference panels, and compared to the truth set to determine the accuracy of imputation.

The nine haplotype reference panels varied in size and composition. Five haplotype reference panels contained 150 cattle (full panels) of which either 0%, 10%, 25%, 50% or 100% were from the BSW breed (*i*.*e*., the breed of the target samples). The other four panels contained either 75 or 30 cattle (reduced panels) that were either from the BSW breed or from breeds other than BSW. DeepVariant identified between 17,035,514 and 28,755,400 sequence variants in the nine haplotype reference panels (Table 2). The full BSW panel contained around 5,167,875 less biallelic SNPs than the full non-BSW panel. The 50% multibreed panel had the highest number of variants shared with the truth set and the lowest number of variants present in the truth set but missing in the reference panel, closely followed by the BSW panel. The reduced non-BSW panel (30 samples) had the lowest number of shared variants and the highest number of variants that were present in the truth but missing in the reference.

**Table 2.** General overview of the haplotype reference panels: number of samples, coverage and number of variants called. Shared and private variants are considered through exact matching (position and alleles). Values are the mean of 3 replicas per haplotype panel.

| Panel | Samples | Coverage | Variants | Biallelic SNPs | SNPs shared truth-query sets | Truth SNPs missing in haplotype panel | SNPs private to haplotype panel |
| --- | --- | --- | --- | --- | --- | --- | --- |
| BSW | 150 | 9.40 | 22,493,568 | 19,682,362 | 13,537,126 | 317,806 | 6,145,236 |
| BSW | 75 | 9.65 | 19,883,488 | 17,345,201 | 13,373,462 | 481,470 | 3,971,739 |
| BSW | 30 | 9.42 | 17,035,514 | 14,839,600 | 12,810,541 | 1,044,391 | 2,029,059 |
| Multibreed (50%) | 150 | 10.48 | 27,710,504 | 24,325,185 | 13,568,744 | 286,188 | 10,756,441 |
| Multibreed (25%) | 150 | 10.86 | 28,755,400 | 25,266,484 | 13,531,721 | 323,211 | 11,734,763 |
| Multibreed (10%) | 150 | 11.44 | 28,608,506 | 25,126,433 | 13,427,451 | 427,481 | 11,698,982 |
| Non-BSW | 150 | 11.78 | 28,303,738 | 24,850,237 | 13,075,827 | 779,105 | 11,774,410 |
| Non-BSW | 75 | 11.78 | 25,059,239 | 21,968,792 | 12,868,909 | 986,023 | 9,099,883 |
| Non-BSW | 30 | 11.45 | 21,011,311 | 18,402,870 | 12,283,284 | 1,571,648 | 6,119,586 |

### Assessment of lcWGS imputation with the different haplotype panels

Increasing the number of reference haplotypes enabled higher F1, recall and precision scores in all tested scenarios (Figure 4A & Supplementary Table 5). Imputation accuracy also improved with increasing lcWGS coverage, with the largest change between 0.01x and 1x coverage. Accuracy continued to improve with diminishing returns between 1x and 4x coverage. The difference in accuracy between panels also reduced as coverage increased.

**Fig. 4.**
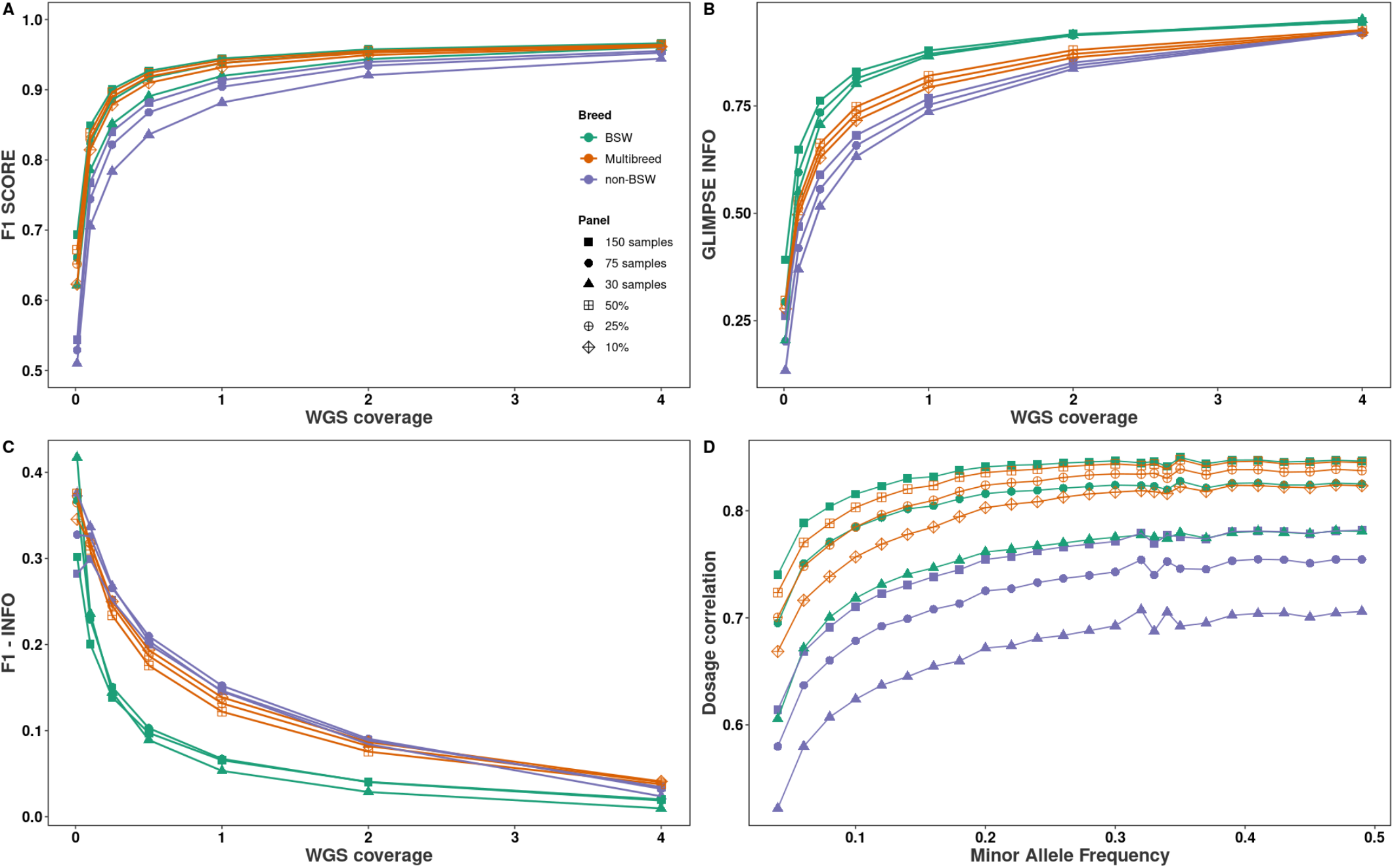
Genotyping accuracy from low-pass whole-genome sequencing. a) F1 score between truth and imputed variants. b) GLIMPSE INFO score achieved with different sequencing coverages and haplotype panels. c) Differences (subtraction) between F1 and GLIMPSE INFO average scores for different sequencing coverages and haplotype panels. d) Squared dosage correlation (r^2^) between imputed data and truth set, stratified by MAF for lcWGS at 0.5x. Panels are indicated with colours and number/percentages of BSW samples in different point shapes. Points indicate the average of the results for all variants in three different replicates.

The largest BSW haplotype reference panel (n = 150) performed better than any of the multibreed panels at all sequencing coverages. Multibreed panels outperformed BSW panels with a larger number of BSW samples, especially at low coverage. For instance, a large multibreed panel containing 10% BSW samples (n = 15) produced higher F1 scores than a smaller breed-specific panel containing two times more BSW samples (n = 30). Similarly, a large multi-breed panel containing 25% BSW samples (n = 37) provided higher F1 scores than a smaller breed-specific panel contain-ing two times more BSW samples (n = 75) for lcWGS below 1-fold coverage. Accuracies were similar between large multibreed panels and smaller breed-specific panels when the coverage of the lcWGS was higher than 1-fold. All results were validated by three different conformations of the haplotype reference panels (replicas). Standard errors accounting for all the replicas did not overlap for any of the haplotype panels (Supplementary Figure 3A).

The imputation accuracy estimated by GLIMPSE (INFO score) was higher for all BSW panels than for the multibreed panels across all coverages (Figure 4B). A higher proportion of variants were imputed with an INFO greater than 0.6 in the BSW than in non-BSW or multibreed panels (Supplementary Figure 3B). Therefore, panels for which the average INFO was higher had also a major proportion of variants with high imputation quality, potentially selected for downstream analyses. Differences between BSW panels and the rest were higher than the differences between multibreed and non-BSW. The average values of F1 and the average INFO scores were closer for the variants imputed with BSW panels (Figure 4C). The differences between both metrics decreased as the coverage of the lcWGS increased (Supplementary Figure 3B-C).

The variants were then stratified by MAF, and the squared correlation of genotype dosages (r^2^) was calculated (Figure 4D). The correlations increased along with the MAF similarly for all the panels. The highest correlations were for BSW panel (150 samples) and multibreed panels (50% and 25%). The values increased substantially between 0-0.1 MAF and continued slowly incrementing until 0.5 for all panels.

## Discussion

Higher F1 scores against a microarray truth set, improved k-mer based variant filtering, and less Mendelian errors suggest that DeepVariant is a superior variant caller to GATK for bovine short read sequencing. These results extend evidence of DeepVariant’s greater accuracy established in multiple human studies (17–20). Ti:Tv ratios in the expected range of 2-2.2 suggest that variant calls private to DeepVariant contain genuine variants, whereas a lower Ti:Tv ratio in vari-ants private to GATK indicate an excess of false positives.DeepVariant revealed more SNPs with impactful annotations, likely providing additional putative trait-associated candidates for downstream analyses. DeepVariant was approximately 3.5x faster in end-to-end variant calling compared to GATK, due to greater multithreading potential and not requiring pre-processing like GATK’s base recalibration step (Supplementary Table 6). The peak memory usage was approximately 65% higher for DeepVariant compared to GATK (81 GB versus 49 GB). Although our work focused on CPU-only machines, DeepVariant also natively offers GPU acceleration (roughly 1.9x faster overall), while GATK has no official GPU support, although there are third-party developments (roughly 1.4x faster overall) (55).

To the best of our knowledge, our study is the first to establish bovine haplotype reference panels with DeepVariant. A within-breed panel consisting of 75 samples enabled us to genotype more than 13 million sequence variants in animals sequenced at 0.5-fold sequencing coverage with F1 scores greater than 0.9. Larger haplotype reference panels (n = 150) from the same breed as the lcWGS data outperform multi-breed panels across all low coverage spectrum (from 0.1- to 1-fold) and MAFs, including rare variants. The development of such panels is a feasible alternative option to using much bigger multibreed panels, such as the 1000 Bull Genomes project imputation reference panel (13). Such large panels, encompassing huge within- and across-breed diversity, may be regarded as the most complete and thus best genomic resources available in bovine genomics. However, using such large panels may be detrimental for breed-specific imputation (also described by Nawaz *et al*. (56)), as we observed many relevant sites were filtered out before imputation due to being multiallelic, resulting in a lower F1 score than the 75 sample BSW panel at 1-fold coverage and greater. Same-breed panels are also more computationally efficient and are 18%-33% faster than using multi- or different-breed panels of the same size (Supplementary Figure 4), and approximately 7 times faster than using the 1000 Bull Genomes Project panel.

In absence of an adequately sized breed-specific panel (*e*.*g*., below 30 animals), F1 scores of 0.9 can also be accomplished either by increasing the coverage of the lcWGS or by adding distantly related samples from other breeds to the haplotype panels as even animals from seemingly unrelated breeds may share short common haplotypes. Both options will provide accurate sequence variant genotypes at affordable costs for samples from rare breeds, where large breed-specific haplotype reference panels can’t be easily established. For instance, F1 scores *>* 0.92 are observed at 2-fold sequencing coverage for all tested haplotype panels with small differences among them. This is likely caused because higher coverages provide more information for imputation from the own sequencing reads, while lower coverages rely on the information from haplotypes in the panels. We also achieved F1 scores of 0.9 with large multibreed panels containing only 10% same-breed samples (n = 15). However, reference panels that contain only few samples from the target breed are in general less informative as evidenced by the lack of around 100K truth SNPs that were present in same-size breed-specific panels. Additionally, a threshold of non-related haplotypes from where only marginal gains to imputation accuracy are observed have been described (15, 56, 57). Overall results are compatible with similar studies with haplotype panels of both bigger and smaller sample sizes (15, 56, 58). Genotypes imputed from lcWGS enable predicting genomic breeding values and facilitate powerful genome-wide association studies at nucleotide resolution (3, 59).

Although imputation accuracy (F1) and GLIMPSE’s predicted imputation accuracy (INFO score) are respectively averaged over each sample and each variant, we note that F1 (truth) is strictly higher than INFO (estimation). The differences appear to be more pronounced for reference haplotype panels that are of different breed to the target sample and at lower coverages (*i*.*e*., less than 0.25-fold coverage, where GLIMPSE’s INFO scores are inaccurate (6)). While, for example, multibreed panels are near equally accurate to the 150 sample BSW panel, the INFO scores are noticeably lower. Similarly, the INFO score drops more rapidly for lower coverages, suggesting that a fixed threshold may be unnecessarily conservative given the slower decay in F1. The GLIMPSE INFO score is also positively correlated with variant MAF, and thus filtering based on INFO predominantly removes low-frequency variants. While INFO and other imputation accuracy scores are still useful, additional care should be taken in determining a constant filtering threshold as more and different panels become available for use.

## ACKNOWLEDGEMENTS & FUNDING

The authors acknowledge the Functional Genomics Center Zürich for generating DNA sequencing data.

This work was supported by grants from the Swiss National Science Foundation (310030 185229) and the European Union’s Horizon 2020 research and innovation programme under Grant Agreement No. 815668 (BovReg).

