## Supplementary Figures for "Size and composition of haplotype reference panels impact the accuracy of imputation from low-pass sequencing in cattle"

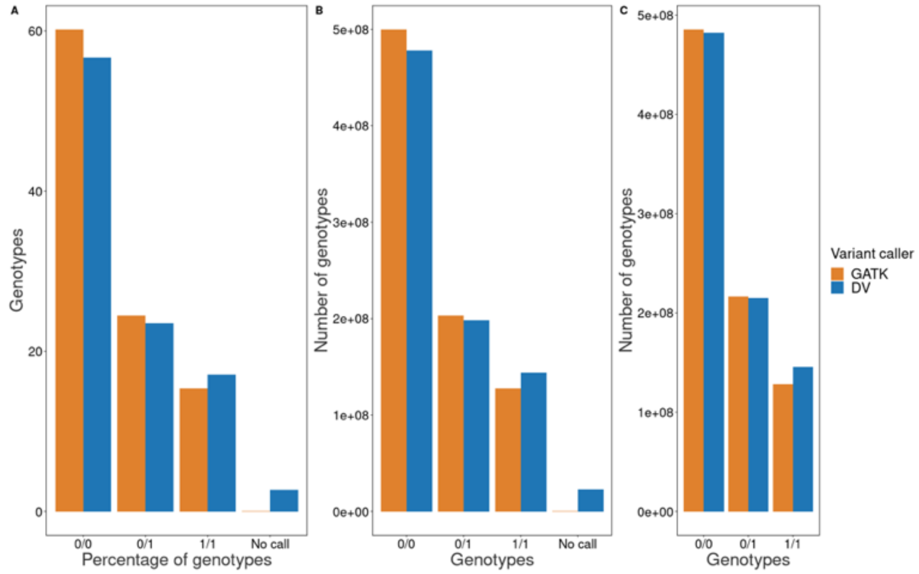


***Figure 1***. *Summary of genotypes. a) Percentage of filtered genotypes called by each variant caller. b) Number of filtered genotypes called by each variant caller. d) Number of imputed genotypes called by each variant caller.*

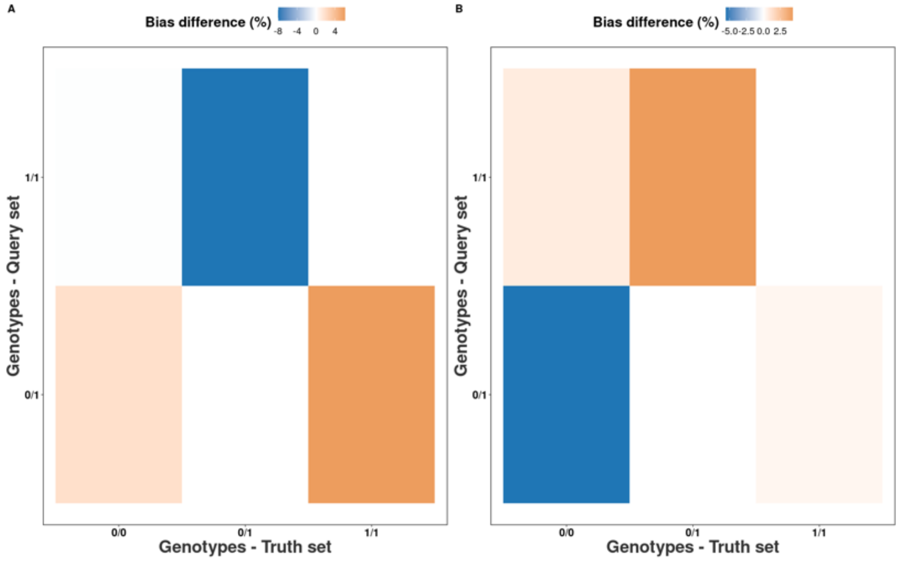


***Figure 2***. *Genotyping accuracy of variant calls. Categorisation and comparison of filtered (a) and imputed (b) genotypes detected with hap.py, where the colour intensity indicates the percentual differences between GATK and DeepVariant (DV).*

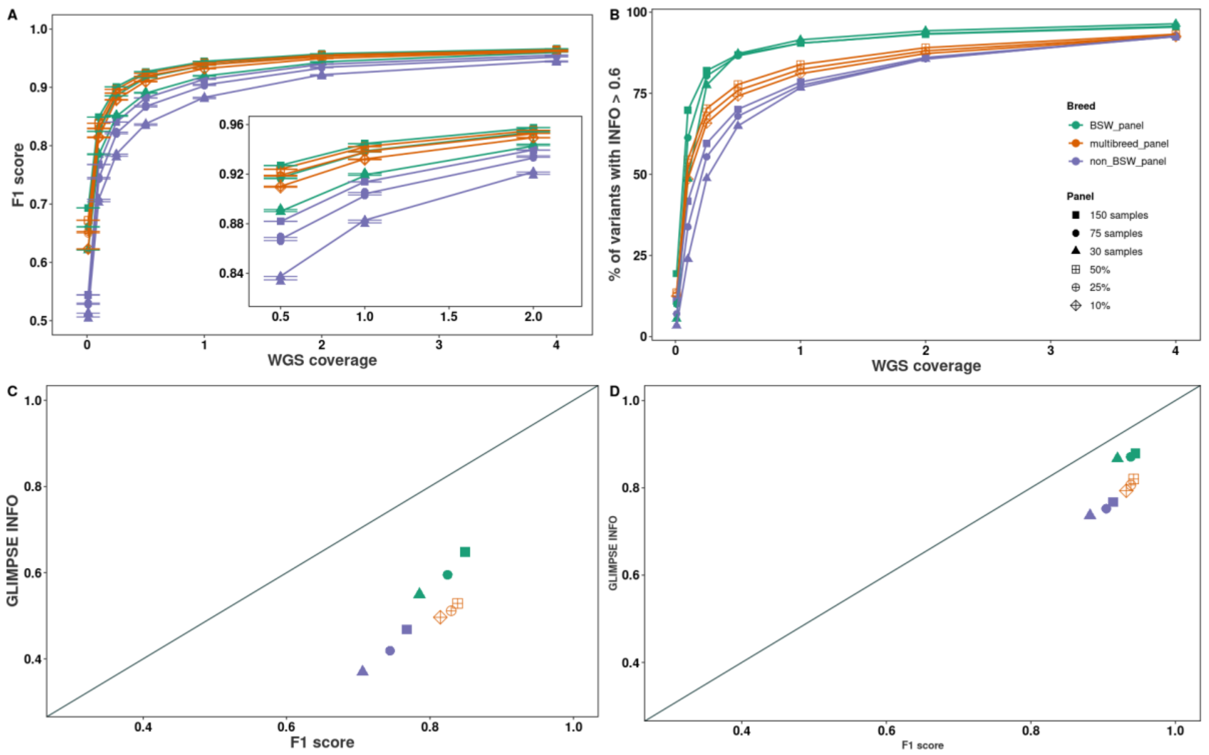


***Figure 3.*** *Genotyping accuracy from low-pass whole-genome sequencing. a) F1 score between truth and imputed variants, with error bars accounting for the three replicas. b) Percentage of imputed variants with a GLIMPSE's estimated accuracy higher than 0.6. Relationship between F1 score and estimated imputation accuracy for lcWGS at 0.1x (c) and 1x (d). Panels are indicated with colours and number/percentage of BSW samples in different point shapes.*

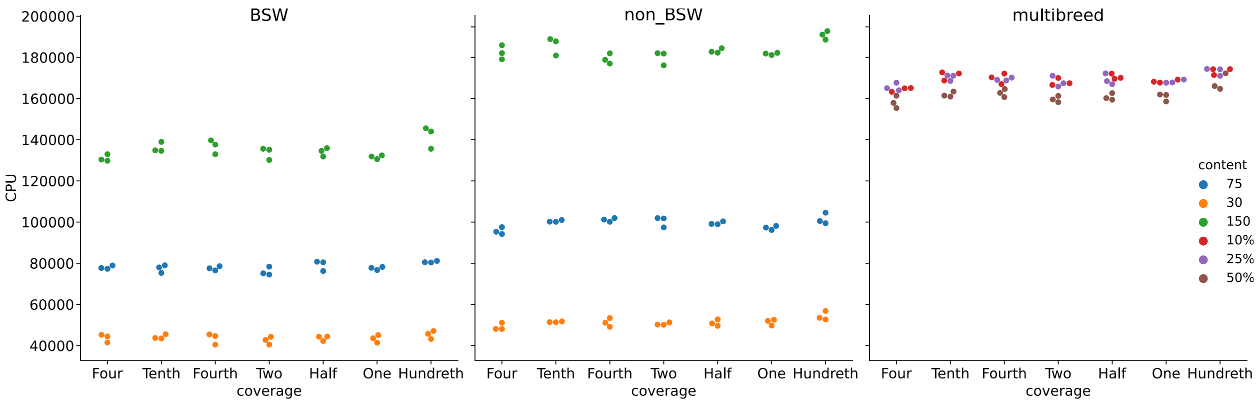


***Figure 4.*** *CPU hours required to impute different coverages and panels for 3 replicates. Compute time was dominated by panel size followed by panel composition, while the input coverage had limited effect.*
