## Supplementary Tables for "Size and composition of haplotype reference panels impact the accuracy of imputation from low-pass sequencing in cattle"

***Table 1.*** *Percentage of overlapping variants across the different GATK and DV sets. Two different intersection modes were used: exact match (same coordinates, and REF and ALT alleles) and position match (only coordinates were queried - in parentheses).*

| | | Overlapping Variants (%) | Overlapping SNPs (%) | Overlapping INDELs (%) |
|---|---|---|---|---|
| **Biallelic** | **GATK** | 93.39 (93.55) | 95.37 (95.38) | 77.97 (79.35) |
| | **DV** | 91.98 (92.15) | 92.61 (92.71) | 86.47 (87.18) |
| **Multiallelic** | **GATK** | 37.17 (60.45) | 24.73 (25.82) | 41.26 (71.84) |
| | **DV** | 38.10 (61.95) | 17.52 (27.94) | 34.41 (60.02) |

***Table 2****. Number of total and multiallelic SNPs shared and private for the different GATK and DV sets. Two different intersection modes were used: exact match (same REF and ALT alleles) and position match (only coordinates were queried - in parentheses).*

| | Overlapping variants | | | Private variants | | |
|---|---|---|---|---|---|---|
| | **SNPs** | **Multiallelic SNPs** | **Multiallelic SNPs (%)** | **SNPs** | **Multiallelic SNPs** | **Multiallelic SNPs (%)** |
| **GATK** | 14,071,834 (14,134,549) | 36,257 (45,367) | 0.26 (0.32) | 791,774 (729,059) | 13,215 (4,105) | 1.67 (0.56) |
| **DV** | 14,071,834 (14,244,141) | 36,257 (39,134) | 0.26 (0.27) | 1,289,951 (1,117,644) | 6,642 (3,765) | 0.51 (0.34) |

***Table 3****. Biallelic variants (SNPs / INDELs) annotated with VEP and classified depending on the likely functional effects: high, moderate, low and modifier. Variants were considered before and after filtering (exact match for filtered out). Variants were also divided depending on whether were called by both variant callers (shared) or only one (private) – position matches were accepted.*

| | Set | High | Moderate | Low | Modifier |
|---|---|---|---|---|---|
| **GATK** | Raw | 2,680 / 4,493 | 67,180 / 1,657 | 90,955 / 3,403 | 15,667,553 / 1,985,337 |
| | Raw private | 546 / 2,283 | 10,390 / 674 | 11,327 / 990 | 926,538 / 452,647 |
| | Raw shared | 2,134 / 2,210 | 56,790 / 983 | 79,628 / 2,413 | 14,741,015 / 1,532,690 |
| | Filtered | 2,252 / 3,993 | 57,675 / 1,522 | 82,618 / 3,126 | 14,574,453 / 1,883,261 |
| | Filtered private | 362 / 2,089 | 7,503 / 598 | 8,714 / 820 | 664,190 / 401,763 |
| | Filtered shared | 1,890 / 1,904 | 50,172 / 924 | 73,904 / 2,306 | 13,910,263 / 1,481,498 |
| | Filtered out | 428 / 500 | 9,505 / 135 | 8,337 / 277 | 1,093,100 / 102,076 |
| **DV** | Raw | 3,530 / 2,778 | 69,525 / 1,162 | 88,839 / 2,833 | 16,125,916 / 1,748,769 |
| | Raw private | 1,396 / 572 | 12,733 / 175 | 9,213 / 421 | 1,384,901 / 216,078 |

| | | | | | |
|---|---|---|---|---|---|
| | Raw shared | 2,134 / 2,206 | 56,792 / 987 | 79,628 / 2,412 | 14,741,015 / 1,532,691 |
| | Filtered | 2,474 / 2,240 | 58,457 / 1,068 | 80,765 / 2,730 | 15,013,146 / 1,700,025 |
| | Filtered private | 584 / 338 | 8,283 / 142 | 6,863 / 424 | 1,102,883 / 218,527 |
| | Filtered shared | 1,890 / 1,902 | 50,174 / 926 | 73,902 / 2,306 | 13,910,263 / 1,481,498 |
| | Filtered out | 1,061 / 612 | 11,214 / 135 | 8,226 / 348 | 1,142,970 / 134,643 |

***Table 4***. *VEP annotation of GATK and DV private variants. MAFs greater than 0.05 are bolded, and HOMALT samples refers to the number of samples with 1/1 genotypes for the variant (out of a maximum of 33 genotyped samples). Some variants were present in the other caller but as a different variant than predicted by truth. Truly missing variant calls are indicated by "-".*

| Variant caller | Position | Predicted impact | MAF | HOMALT samples | Reason not intersecting |
|---|---|---|---|---|---|
| GATK | 9:78221981 | Moderate | 0.78 | 21 | INDEL |
| GATK | 19:46232626 | High | 0.02 | 0 | - |
| DV | 5:116965831 | Low | 0.09 | 1 | - |
| DV | 6:37374718 | Low | 1 | 33 | - |
| DV | 11:15257933 | Low | 0.57 | 10 | INDEL |
| DV | 15:1094876 | Low | 0.75 | 16 | - |
| DV | 25:20777430 | Low | 0.13 | 1 | - |
| DV | 15:49709135 | Moderate | 0.06 | 2 | - |
| DV | 18:61285359 | Moderate | 0.08 | 0 | Multiallelic |
| DV | 4:49174950 | High | 0.02 | 0 | INDEL |
| DV | 7:41125658 | High | 0.26 | 1 | Multiallelic |
| DV | 10:27971463 | High | 0.01 | 0 | INDEL |
| DV | 15:49811331 | High | 0.29 | 3 | - |

***Table 5***. *F1, recall and precision scores when comparing the truth set and the query sets (different coverages and panels). Numbers in pure panels indicate the number of samples. Percentages in multibreed panels indicate the proportion of BSW samples. The top values for each metrics and coverage are highlighted.*

| Metric | Panel | 4x | 2x | 1x | 0.5x | 0.25x | 0.1x | 0.01x |
|---|---|---|---|---|---|---|---|---|
| **F1** | **BSW (150)** | **0.9663** | **0.9575** | **0.9456** | **0.9269** | **0.9009** | **0.8490** | **0.6934** |
| | **BSW (75)** | 0.9655 | 0.9543 | 0.9381 | 0.9167 | 0.8855 | 0.8245 | 0.6609 |
| | **BSW (30)** | 0.9608 | 0.9436 | 0.9198 | 0.8906 | 0.8513 | 0.7853 | 0.6215 |
| | **Multibreed (50%)** | 0.9642 | 0.9554 | 0.9422 | 0.9239 | 0.8966 | 0.8385 | 0.6722 |
| | **Multibreed (25%)** | 0.9629 | 0.9529 | 0.9382 | 0.9188 | 0.8901 | 0.8296 | 0.6518 |

| | | | | | | | | |
|---|---|---|---|---|---|---|---|---|
| | **Multibreed (10%)** | 0.9614 | 0.9494 | 0.9320 | 0.9099 | 0.8786 | 0.8144 | 0.6229 |
| | **Non-BSW (150)** | 0.9552 | 0.9394 | 0.9138 | 0.8819 | 0.8405 | 0.7677 | 0.5442 |
| | **Non-BSW (75)** | 0.9526 | 0.9342 | 0.9042 | 0.8676 | 0.8218 | 0.7443 | 0.5291 |
| | **Non-BSW (30)** | 0.9444 | 0.9208 | 0.8818 | 0.8361 | 0.7837 | 0.7061 | 0.5097 |
| **Recall** | **BSW (150)** | **0.9700** | **0.9565** | **0.9380** | **0.9144** | **0.8810** | **0.8167** | **0.6423** |
| | **BSW (75)** | 0.9677 | 0.9515 | 0.9293 | 0.9008 | 0.8607 | 0.7858 | 0.6064 |
| | **BSW (30)** | 0.9591 | 0.9367 | 0.9059 | 0.8686 | 0.8189 | 0.7394 | 0.5679 |
| | **Multibreed (50%)** | 0.9665 | 0.9528 | 0.9338 | 0.9092 | 0.8736 | 0.8018 | 0.6170 |
| | **Multibreed (25%)** | 0.9650 | 0.9500 | 0.9293 | 0.9034 | 0.8665 | 0.7921 | 0.5975 |
| | **Multibreed (10%)** | 0.9628 | 0.9459 | 0.9222 | 0.8934 | 0.8535 | 0.7756 | 0.5702 |
| | **Non-BSW (150)** | 0.9523 | 0.9317 | 0.8993 | 0.8596 | 0.8100 | 0.7243 | 0.4961 |
| | **Non-BSW (75)** | 0.9457 | 0.9222 | 0.8849 | 0.8394 | 0.7835 | 0.6939 | 0.4760 |
| | **Non-BSW (30)** | 0.9282 | 0.8995 | 0.8526 | 0.7974 | 0.7350 | 0.6475 | 0.4535 |
| **Precision** | **BSW (150)** | 0.9626 | **0.9586** | **0.9512** | **0.9398** | **0.9218** | **0.8839** | **0.7534** |
| | **BSW (75)** | **0.9632** | 0.9571 | 0.9472 | 0.9331 | 0.9118 | 0.8672 | 0.7262 |
| | **BSW (30)** | 0.9625 | 0.9507 | 0.9340 | 0.9138 | 0.8863 | 0.8373 | 0.6862 |
| | **Multibreed (50%)** | 0.9620 | 0.9580 | 0.9507 | 0.9392 | 0.9208 | 0.8788 | 0.7383 |
| | **Multibreed (25%)** | 0.9609 | 0.9559 | 0.9473 | 0.9346 | 0.9152 | 0.8709 | 0.7171 |
| | **Multibreed (10%)** | 0.9599 | 0.9530 | 0.9419 | 0.9269 | 0.9051 | 0.8572 | 0.6864 |
| | **Non-BSW (150)** | 0.9583 | 0.9473 | 0.9287 | 0.9054 | 0.8746 | 0.8167 | 0.6027 |
| | **Non-BSW (75)** | 0.9597 | 0.9465 | 0.9245 | 0.8978 | 0.8641 | 0.8026 | 0.5956 |
| | **Non-BSW (30)** | 0.9611 | 0.9431 | 0.9133 | 0.8788 | 0.8392 | 0.7763 | 0.5818 |

***Table 6***. *Compute resources used by DeepVariant (DV) and GATK to pre-process aligned BAM files, call variants per sample (gVCF stage), and jointly genotype and filter variants (pVCF stage). Times are listed as CPU hours, and peak memory usage across all stages is given in gigabytes. DV does not require pre-processing. Jobs were submitted to nodes with different CPUs and non-exclusive use, and so these figures do not represent precise benchmarking. However, any node/usage variability is minor compared to the differences in total CPU hours used between DV and GATK.*

| Variant caller | Pre-processing (h) | gVCF stage (h) | pVCF stage (h) | Peak memory (GB) |
|---|---|---|---|---|
| DV | - | 577 | 3 | 81 |
| GATK | 820 | 999 | 233 | 49 |
